# Temporal analysis of reproduction distributed in space illuminates the climate-change resiliency of toyon (*Heteromeles arbutifolia*)

**DOI:** 10.1101/2025.09.09.675207

**Authors:** Daniel Dakduk, Jeremy B. Yoder

## Abstract

**Premise:** Toyon, *Heteromeles arbutifolia* (Lindl.) M. Roem. (Rosaceae), is an iconic and ecologically important member of California chaparral and oak woodland communities. Toyon’s habitat faces changing wildfire regimes, widening variation in annual rainfall, and competition by introduced species. We used a new modeling method, temporal analysis of reproduction distributed in space (TARDIS) to examine how recent climate change alters habitat suitability for toyon.

**Methods:** As data for TARDIS, we annotated flowering and fruiting in images from 4,105 observations of toyon contributed to the iNaturalist crowdsourcing platform. From these records we trained Bayesian additive regression tree models relating weather to toyon flowering. We used a trained model to hindcast flowering each year back to 1900, and examined trends in hindcast flowering. For comparison, we also modeled changing habitat suitability using a conventional species distribution model (SDM) relating toyon presence to 30-year climate averages.

**Key results:** Toyon flowering is associated with greater winter precipitation and warmer fall and winter temperatures. Our hindcast finds mean flowering intensity has been stable to slightly increasing since 1900, with greater increases at higher elevations, but also at lower latitudes. Variation in flowering intensity has also increased, especially at lower latitudes. Trends in flowering are positively correlated with changes in SDM-predicted suitability.

**Conclusions:** TARDIS recovers biologically realistic predictors of toyon flowering, and hindcast changes in flowering intensity indicate the species’ range remains suitable after 125 years of changing climate. Overall our results indicate toyon populations remain healthy, but may have limited opportunity to migrate northwards as climate change continues.

## Introduction

Species distributions are complex phenomena, influenced by a variety of biotic and abiotic factors. Spatial variation in climate, resources, competitors, and mutualists interact with a species’ needs and tolerances to define where they can establish self-sustaining populations (Hutchinson, 1957; MacArthur, 1972). These factors directly shape how species geographic distributions may change in response to global climate change (Morelli et al., 2016; Pulliam, 2000). Changing climates have driven plant populations to higher, and cooler, latitudes and elevations, resulting in reduced geographic ranges, changing community compositions, and disrupted interactions with other species (Gómez-Ruiz and Lacher, 2019; Kelly and Goulden, 2008; Telwala et al., 2013; Wróblewska and Mirski, 2018; Zu et al., 2021). Anticipating and mitigating the impacts of global climate change on biodiversity requires better understanding of how geographic distributions are shaped by species’ physiological tolerances, resource needs, and interactions with other taxa.

Species distribution models (SDMs) have long been a useful way to map geographic distributions for species of interest, identify environmental factors that determine species’ presence on the landscape, and project how that distribution may change under future conditions (Elith and Leathwick, 2009; Guisan and Zimmermann, 2000). SDMs are usually one of a variety of different machine-learning algorithms, which in their simplest form pair presence/absence records for a species with environmental data for those locations (Elith and Leathwick, 2009; *Merow et al., 2014b*). This allows SDMs to estimate the probability of a species’ occurrence in locations and times where it has not been directly observed (Guisan and Zimmermann, 2000). SDMs have been used to estimate current geographic distributions for species of interest, and analyze the overlaps in those ranges (Esque et al., 2023; Godsoe et al., 2009). They have also been used to identify future suitable habitat under projected future climate scenarios (Carlson et al., 2022; Hamann and Wang, 2006; Shryock et al., 2025).

Most SDMs relate environmental variables to a species’ presence or absence on the landscape to make projections of its distribution (Guisan and Zimmermann, 2000; Merow et al., 2014*b*). SDMs may also use “mechanistic” methods, defining mathematical functions based on a species’ demographic status in different environments, or the relationships between environmental variables and its physiological tolerances or functional traits (Dormann et al., 2012; Merow et al., 2014a). Rather than correlating the presence of a species with potentially predictive environmental variables, mechanistic models link environmental variables with specific traits of an organism, such as physiology, and then project this information onto the landscape (Dormann et al., 2012). This is particularly useful for determining the range limits of a species, because it allows researchers to directly test how specific environmental factors limit a species’ population dynamics, and thereby geographic distribution (Kearney and Porter, 2009). Mechanistic modeling has been used to predict life-cycle completion in invasive common ragweed [*Ambrosia artemisiifolia* L. (Asteraceae)] and determine its prospective habitat in Europe (Chapman et al., 2017), and to map the spread of Oriental bittersweet [*Celastrus orbiculatus* Thunb. (Celastraceae); Merow et al. 2011] in North America, among other applications. Mechanistic modeling requires more intensive data collection to characterize physiological tolerances or demographic status, or both, in different environments (Dormann et al., 2012; Merow et al., 2014*b*). Notably, both mechanistic and “correlative” (presence-absence) SDMs can be treated as hypotheses about what factors define a species’ geographic distribution, by testing model predictions against independent observations of the species’ presence, absence, demographic status, or physiological condition (Radomski, 2025).

A possible middle ground between presence-based SDMs and mechanistic models is to model the presence or intensity of demographic processes, such as reproductive effort or recruitment, across years. This approach lets us evaluate annual variation in population activity relative to annual variation in weather or other environmental factors across a species’ range, providing something closer to the information in a mechanistic model of population growth, but with data that is more accessible for many species. A recent study of Joshua trees [*Yucca brevifolia* Engelm. and *Y. jaegeriana* McKelvey ex. Lenz (Asparagaceae)] used geographically dispersed records of the trees’ flowering and fruiting over 15 years to model weather conditions associated with flowering (Yoder et al., 2024). The trained model could hind-cast past Joshua tree flowering events from historic weather records, and Yoder *et al* (2024) used the hindcast to identify regions of the species’ current range where conditions favoring flowering have increased in frequency, and decreased, as a result of recent climate trends. This approach, which we now call temporal analysis of reproduction distributed in space (TARDIS), works from records that can be casually collected — Yoder *et al*. (2024) used annotated, validated records contributed to the iNaturalist crowdsourcing platform (inaturalist.org) — and which are available for many other plant species.

In this study, we adapted the techniques developed by Yoder *et al*. (2024) to model flowering activity in toyon [Fig. 1, *Heteromeles arbutifolia* (Lindl.) M. Roem. (Rosaceae)]. Toyon is largely endemic to the California Floristic Province (CFP), distributed along the coast and inland to the western slopes of the Sierra Nevadas (Fig. 1E; Montalvo et al. 2018). Toyon is an ecologically important, often dominant shrub in chaparral and oak woodland communities, and it has been an important source of food, medicine, and wood for Native Americans living in its range (Higgins, 2019; Incayawar, 2010). The name *toyon* is a loan word from the Ohlone people, in the north of the species’ range (Higgins, 2019); the Tongva people farther south call it *ashuwet* (Incayawar, 2010). Toyon’s evergreen leaves and bright red, berry-like pomes have made it popular for landscaping (Montalvo et al., 2018), and inspired a folk history supposing it to be the “holly” from which Hollywood takes its name — though the actual historical record contradicts this (Higgins, 2019).

**Fig. 1.**
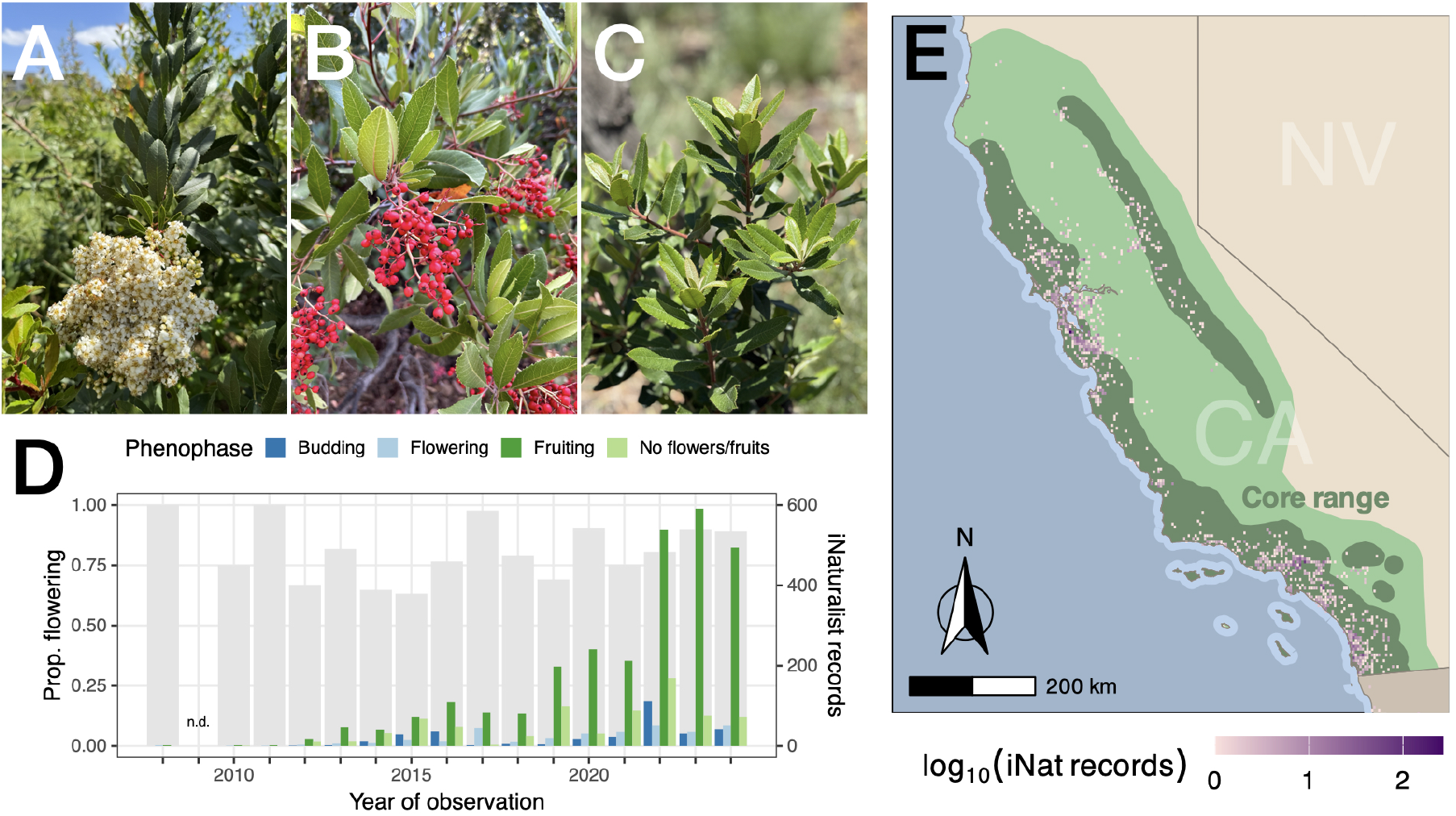
Observations of toyon reproductive activity contributed to iNaturalist. (A-C) Images of toyon with flowers, fruits, and no evidence of flowering (from iNaturalist records 281184791 by user ‘ahalligan’; 282517568 by user ‘kd_ods’; and 276841543 by user ‘ecowriter’); (D) Counts of research-grade phenophase-annotated records from 2008– 2024, by year and phenophase annotation (colored bars, right axis) compared to the proportion of records in each year indicating flowering (i.e., with buds, flowers, or fruit; gray bars, left axis); (E) Distribution of iNaturalist records relative to toyon’s core range in California, USA (pink to purple shading, *log*_10_ of records within 4km grid cells; dark green shading indicates the core range (U.S. Geological Survey, 1999), lighter green shading indicates our designated study area)

Toyon is not considered endangered (IUCN SSC Global Tree Specialist Group & Botanic Gardens Conservation International (BGCI), 2020), but has considerable conservation value as a foundational species for habitat restoration (Riordan et al., 2018). Its range includes regions expected to lose biodiversity under warmer future climate scenarios, and which face increasing frequency and severity of wildfire (Harrison et al., 2024; Park et al., 2018). Understanding how toyon’s range is limited by the varying climates it encounters in the complex topography of the CFP, and how that geographic distribution may shift with global climate change, can help land managers and policymakers plan for the protection of living communities in this global biodiversity hotspot.

The Mediterranean climate of the CFP presents toyon populations with hot, dry summers relieved by winter precipitation and, along the coast, fog that persists into the dry season (Vasey et al., 2014). Toyon typically flowers from the end of the wet season in May or June into August, with fruits persisting on the plants into the following year (Montalvo et al., 2018; Phipps, 2012). It has evergreen, sclerophyllous leaves and deep roots (Pezner et al., 2020), and it has been categorized as drought tolerant (Davis and Mooney, 1986; Valladares and Pearcy, 1997). However, toyon seedlings have lower photochemical efficiency and stomatal conductance in full sunlight, and stressful summer conditions in its typical habitat likely limit seedling survival (Valladares and Pearcy, 1997). Possibly as a consequence, toyon is more abundant on deeper, more mesic soils (Meentemeyer et al., 2001). Toyon populations also show some evidence of locally adapted, or plastic, variation in response to climate. Populations in the warmer, drier southern end of the range maintain greater rates of photosynthesis at high temperatures and low water potentials, compared to those found further north (Mooney et al., 1975). Toyon populations at higher elevations also invest more in seed provisioning, though there is no correlation between seed mass and either latitude or annual precipitation (Martijen and Bullock, 1997). Finally, leaf morphology variation is correlated with fitness measures and environmental variation, and shows evidence of both local adaptation and plastic responses to growing environment in a recent greenhouse and common-garden study (Thomas and Prunier, 2024).

Overall, it seems likely that toyon populations, and toyon’s geographic distribution, are limited by climate factors that are shifting with global trends. If this is the case, we would expect to see demographic evidence of population declines in regions that have become less suitable for toyon since the early 20th century, and increasing population growth in regions that have become newly suitable, or which are prospective climate refugia. A comparison of California plant community surveys taken in 1935 and 2005 found that toyon’s optimal elevation had shifted just over 53 meters down-hill, tracking changes in drought stress (Crimmins et al., 2011); one prior SDM study predicts that most of the species current habitat in southern California will remain suitable under future climate scenarios for the mid 21st century (Riordan et al., 2018). Still, no range-wide assessment has examined how toyon populations have experienced climate change so far, which has resulted in an average increase in annual mean temperature of about 1°C across the species’ range (annual mean temperature for 1991-2020 compared to 1901-1930; PRISM Climate Group, Oregon State University 2014).

Here, we applied TARDIS to examine how climate trends since the beginning of the 20th century may have altered the geographic extent of suitable conditions for toyon. Following Yoder *et al*. (2024), we used records of toyon flowering from the iNaturalist crowdsourcing platform to train machine-learning models predicting annual flowering activity with annual variation in weather. We then used a trained model to hindcast toyon flowering activity from weather records for each year from 1900 to 2024, and examined how hindcast flowering has changed over this 125-year study period. Specifically, we tested for increasing flowering activity in regions that are likely climate refugia, either *a priori* — at higher elevations and latitudes — or as identified by a presence-based SDM. We find that toyon’s current range continues to support reproductively active populations, but our results also suggest that this ecologically important species may have limited opportunity to shift its range in response to future climate change.

## Methods

We conducted all analyses in R (version 4.5, R Core Team 2025), with specialized functions as indicated.

### Annotation and compilation of crowdsourced records

We obtained records of *H. arbutifolia* flowering activity from iNaturalist (inaturalist.org), a community science platform that allows contributors to upload photos of wildlife along with information about where the observation was located and when it was observed (iNaturalist community, 2025). We searched iNaturalist for records of *H. arbutifolia* that needed confirmation of species identification, included photos, and were not annotated for plant phenology. We confirmed that the records were correctly identified as *H. arbutifolia* and used annotation tools in the iNaturalist platform to mark the records as showing flowers budding, flowers open, and/or fruit, or showing no evidence of flowering (Figure 1A-C). We used modifications of functions from the package `rinat` to download records from the iNaturalist database using the platform API. We specifically queried the database for “Research Grade” records, which have species identification confirmed by at least two iNaturalist users with no contradictory identifications, from the years 2008 (when iNaturalist launched) through 2024 (the most recent full year available); and we further filtered downloaded records to remove any marked in the database as captive/cultivated or with location uncertainty greater than 1km.

### Linking flowering records to weather data

We obtained monthly weather records from the PRISM database (PRISM Climate Group, Oregon State University, 2014), which provides spatially interpolated raster-formatted data on temperature (minimum and maximum, in degrees Centigrade), vapor pressure deficit (minimum and maximum, in hPa) and total precipitation (in millimeters) across the contiguous United States from 1895 to the present. (Toyon’s range extends south into Baja California, so our analyses are necessarily restricted to the range north of the US-Mexico border, which is a large majority of the total distribution.) We summarized monthly weather records into quarters (January–March, April–June, July– September, and October–December).

To link flowering activity as observed in the iNaturalist records to weather, we aggregated the iNaturalist records by year and grid cell within the ~4km raster grid of the PRISM data layers. From the records within each grid cell in a given year, we calculated an annual frequency of flowering as the proportion of records indicating flowering (annotated with flowers budding, flowers, or fruit) out of the total records within the cell. (Hereafter, we use “flowering frequency” to refer to the values calculated this way, and “flowering intensity” to mean the biological phenomenon approximated by that flowering frequency — the proportion of individuals flowering in a toyon population in a given year and location.) We matched each cell with an annual frequency of flowering to quarterly weather records for the first quarter of the year in which flowering was observed (Y0Q1, i.e., January–March 2024 for observations in 2024), each quarter of the year prior to flowering (from Y1Q4, or October–December 2023, back to Y1Q3, or July–September 2023).

### Modeling flowering activity

We trained Bayesian additive regression tree (BART) models predicting flowering activity in a given year and location with weather variation in the three quarters (Y0Q1 back through Y1Q1) prior to flowering. BART models are sum-of-tree classification and regression tree machine-learning models, which have notable advantages over other ML methods widely used in ecology. BARTs require minimal parameterization; they provide better quantification of uncertainty than similar methods like boosted regression trees; and they offer a clear means to evaluate predictor importance, via the number of times predictors are used for tree splitting, which helps to avoid over-fitting (Carlson, 2020; Carlson et al., 2022; Chipman et al., 2010).

The prototype TARDIS study, by Yoder *et al*. (2024), used binary-response BART models because the focal species in that study had highly bimodal annual flowering activity characteristic of a masting strategy. We trained a binary-response model of toyon flowering following Yoder *et al*. (2024), but because our toyon data revealed more uniformly high frequency of flowering in every year of observation (Fig. 1D, gray bars), we also explored the use of continuous-response BART models. Continuous-response modeling allows better fitting to variation in frequency between the extremes seen in a masting species. We compared the outcomes of these two modeling approaches to select a working model for downstream prediction and analysis.

We used functions in the `dbarts` (Dorie, 2023) and `embarcadero` (Carlson, 2020) packages to train and evaluate binary-response BART models of flowering activity — i.e., models predicting the presence or absence of flowering activity in a given year. Following the approach in Yoder *et al*. (2024) we reduced flowering activity to a binary by treating years and locations with estimated frequency of flowering equal to 0 as having no flowering, and all other years and locations as having flowering. We identified informative predictors using the diagnostic predictor selection procedure provided by `embarcadero` to identify the strongest predictors, training models with 10, 20, 50, 100, or 200 component trees. Predictors that have consistently higher frequency of inclusion in simpler models (i.e., those with fewer trees) can be identified as the most informative predictors (Chipman et al., 2010). After identifying the most informative predictors, we trained a standard BART model for downstream analysis.

We used functions provided in the `SoftBart` package (Linero, 2022; Linero and Yang, 2018) to train and evaluate continuous-response BART models of flowering activity — i.e., models predicting the frequency of flowering in a given year. In addition to training continuous-response models, `SoftBart` implements a predictor selection method enabled by using a Dirichlet-distributed hyperprior for predictor inclusion in regression tree splits. This means better predictors have higher posterior probabilities of inclusion in a trained model. We performed predictor selection by training models with 10, 20, 50, 100, or 200 component trees, and comparing the posterior probabilities of inclusion for all candidate predictors. As with the process for binary-response BARTs, we inferred that predictors with the highest posterior probability of inclusion in simpler models are most informative. We then trained a final working BART model with those most informative predictors.

We compared performance of the binary- and continuous-response models in terms of their root mean squared error (RMSE), and by considering the biological realism of their estimated predictor partial effects. The continuous-response model performed better by both standards (see Results), and so we retained it as the primary working model for downstream analysis.

### Hindcasting flowering activity

We next used our working model to predict flowering intensity (frequency of records indicating flowering) across the range of toyon for each year from 1900 to 2024. We defined the spatial extent for these analyses using a digitized range polygon for toyon from Ebert’s *Atlas of United States Trees* (U.S. Geological Survey, 1999), which we call the species “core range” within California (Fig. 1E, dark green shading), and a larger “study area” defined by a convex hull around iNaturalist records of toyon occurrence, buffered to 50km (Fig. 1E, lighter green shading). Records of toyon occurrence, and iNaturalist observations of toyon flowering activity, extend outside of the “core range” boundaries — toyon has been reported as far north as Oregon (Wood, 2008) — but occurrence records are at their highest density within the core range polygon, and it thus provides a useful point of reference in our examination of toyon’s geographic distribution.

Within the study area, we hindcast annual variation in flowering intensity by using our finalized continuous-response BART model to predict flowering intensity from PRISM-derived weather records for each year from 1900 to 2024. To describe spatial variation in conditions favoring flowering, examine how change in hindcast flowering activity arises from changes in the weather predictors of flowering, and test whether conditions favoring flowering track climate change refugia, we summarized modeled flowering intensity as the mean and coefficient of variation (CV) of hindcast flowering intensity from 1991 to 2020 (a “recent” period, within the 125-year span of our data), and as the mean and CV of hindcast flowering intensity from 1901 to 1930 (an “early” period).

### Validation from independent records

To validate our hindcast predictions of flowering intensity, we drew on records of toyon flowering collected by the USA-National Phenology Network (USA National Phenology Network, 2025). USA-NPN records include fine-grained surveys of toyon leaf-out and fruit development at 197 unique locations from 2011 through 2024. We estimated flowering intensity from these data, as the proportion of survey observations for a given flowering year that reported fruit. Most sites were surveyed in just one or a few years, but in total the USA-NPN records provided 380 site-year observations of flowering intensity. We matched USA-NPN site-year observations to the hindcast predicted intensity of flowering for the same times and locations, and tested whether hindcast probability of flowering was positively correlated with USA-NPN recorded flowering intensity.

### Comparison to expected and modeled climate– driven range shifts

Finally, for comparison to our TARDIS results, we used a conventional presence-based species distribution model to infer how toyon’s suitable habitat has shifted from the early 20th century to the present day. As climate predictors for the early and present-day time periods, we calculated the 19 standard “BioClim” variables (O’Donnell and Ignizio, 2012) from PRISM monthly data, averaging them over 1991–2020 (for the recent period) and 1901–1930 (for the early 20th century).

As training data for the SDM, we downloaded toyon occurrence records from the Global Biodiversity Information Facility (GBIF) database (gbif.org; GBIF 2025) using utilities in the `rgbif` package, which we filtered to remove records with higher location uncertainty and high-probability captive or cultivated status using utilities in the `CoordinateCleaner` package. We removed records from prior to 1991, the start of the period covered by our “recent” climate data, and rasterized the observations to the grid of the PRISM data layers; grid cells containing cleaned records were the presence points for SDM model training. As pseudo-absence records for model training, we randomly drew an equal number of grid cells within the study area (Fig.1E, light green) that contained none of our cleaned GBIF occurrence records (Barbet-Massin et al., 2012).

We then trained a binary-response BART model with BioClim predictor averages for 1991–2020, with predictors selected using the methods described above. We used the trained model to predict habitat suitability for toyon across the study area, both for 1991–2020 and for 1901–1930, using the BioClim predictor averages we had calculated for each time period. Within the study area, we identified regions as gaining suitability if they were predicted to be suitable in 1991–2020 but not in 1901–1930, and losing suitability if they were not suitable in 1991–2020 but suitable in 1901–1930.

We compared these static SDM predictions of changing habitat suitability to the TARDIS hind-cast of changing flowering intensity by testing for a correlation between the change in SDM-predicted probability of occurrence and the change in mean and CV of hindcast flowering intensity from 1901– 1930 to 1991–2020. We also tested whether the change in mean and CV of hindcast flowering intensity was correlated with latitude and elevation, based on an *a priori* expectation that species’ suitable climate conditions shift further north and uphill with global climate change, all else being equal.

## Results

### Flowering activity records obtained from iNaturalist

Our validation, annotation, and data cleaning obtained 4,105 phenophase-annotated records from iNaturalist, spanning 2008–2024 (Fig. 1D). Records were concentrated in more recent years, reflecting the substantial growth of the iNaturalist contributor community since 2008 (we found no records from 2009 that passed our quality thresholds; in the year with the most records, 2022, we found a total of 870). The percentage of records indicating flowering was high in all years with data, ranging from 63% to 100%, consistent both with a reproductive strategy in which toyon flowers and fruits in every growing year, and with the high visibility of the species’ flowers and fruits to iNaturalist contributors. Records were distributed across the species’ range, but more heavily concentrated around the urban population centers of San Francisco, Los Angeles, and San Diego (Fig. 1E). Rasterized to the grid of our PRISM-derived quarterly weather records, these data gave us 2,345 estimates of flowering frequency in particular years and locations, which provide the basis for model training. (PRISM records cover the continental United States, while toyon’s range extends south into Baja California; our analyses are therefore restricted to the large majority of the species’ range north of the US-Mexico border.)

### BART modeling of annual flowering activity

Predictor selection for the binary-response BART model of annual flowering identified six top predictors (Fig. 2): total precipitation in the third quarter of the year before flowering (PPT Y1Q3) and in the fourth quarter of the year before flowering (PPT Y1Q4); minimum vapor pressure deficit in the fourth-quarter of the year before flowering (VPDmin Y1Q4); maximum VPD in the first quarter of the year of flowering (VPDmax Y0Q1); and minimum temperature in the fourth quarter of the year before flowering (Tmin Y1Q4) and the third quarter of the year before flowering (Tmin Y1Q3). A binary-response BART model trained on our data with those predictors had relatively good classification accuracy, with AUC = 0.73, and RMSE = 0.59.

**Fig. 2.**
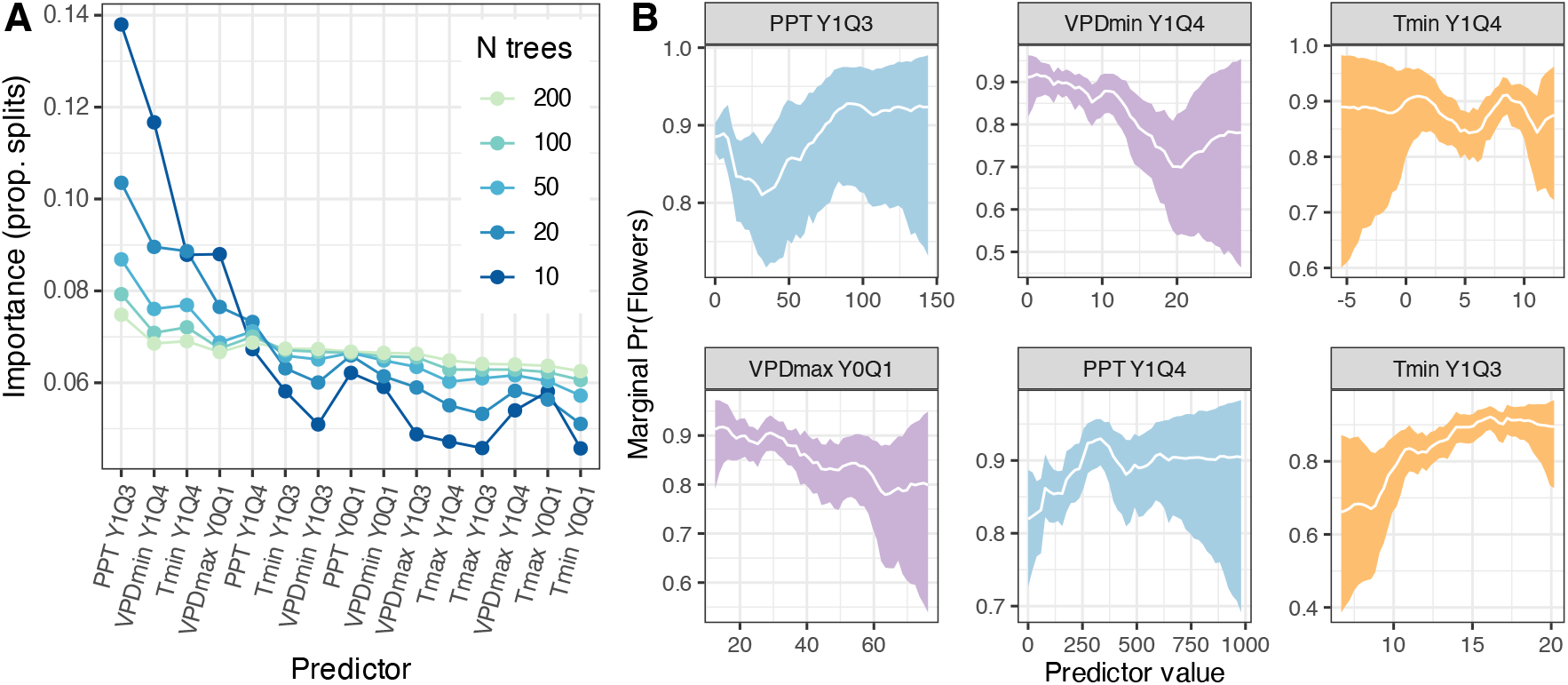
Predictor selection and posterior partial predictor effects for the binary-response BART model of toyon flowering activity. (A) Relative contributions of candidate predictors in models with decreasing complexity (regression tree count) identifies the six left-most predictors as meaningfully predicting flowering activity (Carlson et al., 2022). (B) Partial effects of the top six environmental predictors in the final trained model (white line, median; shaded area 95% density interval across trees, shading color indicates predictor type).

Predictor selection for the continuous-response BART model of annual flowering intensity identified an overlapping but not identical set of six top predictors (Fig. 3): minimum temperature in Y1Q3 and Y1Q4; total precipitation in Y0Q1, Y1Q4, and Y1Q3; and minimum VPD in Y0Q1. The RMSE for a continuous-response model trained with these predictors was 0.35 (classification accuracy is not relevant for a continuous-response model).

**Fig. 3.**
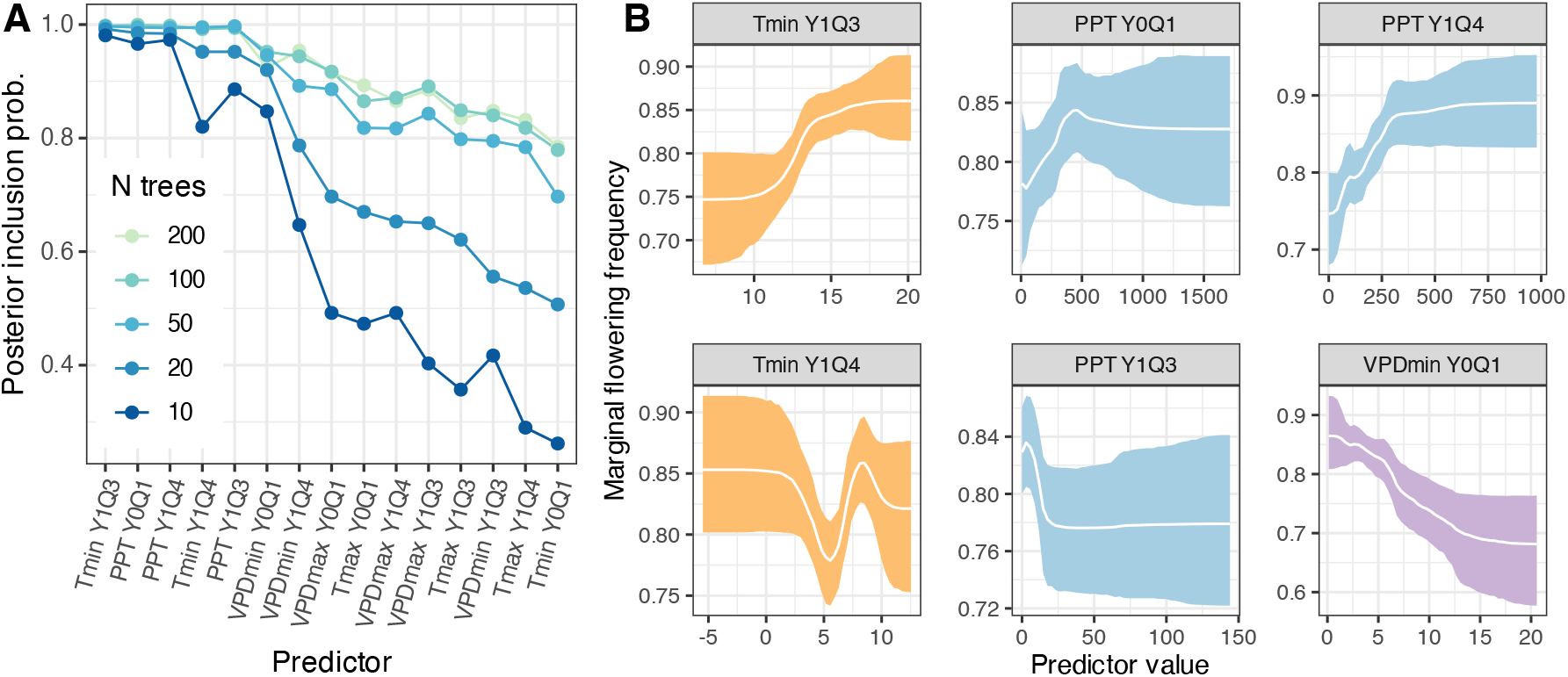
Predictor selection and posterior partial predictor effects for the continuous-response BART model of toyon flowering activity. (A) Posterior inclusion probability for candidate predictors in models with decreasing complexity (regression tree count) identifies the six left-most predictors as meaningfully predicting flowering activity. (B) Partial effects of the top six environmental predictors in the final trained model (white line, median; shaded area 95% density interval across trees, shading color indicates predictor type).

For the four predictors shared between the binary- and continuous-response models (Fig. 2, 3: PPT Y1Q3 and Y1Q4, Tmin Y1Q3 and Y1Q4, and VPDmin Y0Q1) the models estimated qualitatively similar partial effects. Both models found greater flowering intensity associated with greater total precipitation in Y1Q4, with lower minimum VPD in Y0Q1, and with higher minimum temperatures in Y1Q3; they also both found complex, though not entirely similar, relationships between flowering and precipitation in Y1Q3. Given that the two models find broadly similar, biologically realistic relationships between weather and flowering, but the continuous-response model has lower RMSE, we elected to use the continuous-response model for our hindcast of toyon flowering activity and all subsequent analysis.

### Hindcast flowering activity

The hindcast from our continuous-response model of toyon flowering predicts that annual weather variation from 1900 through 2024 is consistent with high frequency of flowering in most years (Fig. 4A,B; Table 1). Across the study area, the median hindcast flowering intensity, averaged over 1991–2020 was 0.84 (95% density from 0.70 to 0.90), and results were similar inside and outside the core range. Inter-annual variation was low, with a median CV of hindcast flowering intensity of 0.06 (95% density from 0.03 to 0.13) across the full study area, though this was somewhat higher within the core range (0.07; 0.03 to 0.14) than outside it (0.05; 0.03 to 0.11).

**Table 1.** Summaries of mean and CV of hindcast flowering intensity over 1901–1930 and 1991–2020, change in mean and CV of hindcast flowering intensity between those periods, and correlations with latitude, elevation, and SDM-predicted probability of occurrence.

|  |  | Correlation* with |  |  |  |
| --- | --- | --- | --- | --- | --- |
|  | Median (95% density) | Latitude | Elevation | SDM Pr(present) | ΔPr(present) |
| *Mean flowering intensity, 1901–1930* |  |  |  |  |  |
| Full study area | 0.80 (0.66, 0.89) | 0.84 | n.s. | 0.04 | – |
| Core range | 0.79 (0.67, 0.88) | 0.81 | 0.07 | -0.19 | – |
| Outside core range | 0.81 (0.66, 0.88) | 0.83 | n.s. | 0.36 | – |
| *Mean flowering intensity, 1991–2020* |  |  |  |  |  |
| Full study area | 0.84 (0.70, 0.90) | 0.84 | 0.03 | -0.05 | – |
| Core range | 0.82 (0.70, 0.91) | 0.82 | 0.05 | -0.20 | – |
| Outside core range | 0.84 (0.69, 0.90) | 0.84 | n.s. | 0.19 | – |
| *Change in mean flowering intensity, 1991–2020 versus 1901–1930* |  |  |  |  |  |
| Full study area | 0.02 (-0.02, 0.08)<sup>†</sup> | -0.22 | 0.04 | – | 0.16 |
| Core range | 0.02 (-0.03, 0.08)<sup>†</sup> | -0.08 | n.s. | – | 0.09 |
| Outside core range | 0.03 (-0.02, 0.07)<sup>†</sup> | -0.34 | 0.07 | – | 0.22 |
| *CV of flowering intensity, 1901–1930* |  |  |  |  |  |
| Full study area | 0.06 (0.02, 0.10) | -0.61 | -0.08 | 0.11 | – |
| Core range | 0.06 (0.02, 0.10) | -0.64 | -0.09 | 0.23 | – |
| Outside core range | 0.05 (0.01, 0.10) | -0.56 | -0.10 | -0.07 | – |
| *CV of flowering intensity, 1991–2020* |  |  |  |  |  |
| Full study area | 0.06 (0.03, 0.13) | -0.63 | -0.10 | 0.21 | – |
| Core range | 0.07 (0.03, 0.14) | -0.72 | -0.09 | n.s. | – |
| Outside core range | 0.05 (0.03, 0.11) | -0.52 | -0.12 | 0.12 | – |
| *Change in CV of flowering intensity, 1991–2020 versus 1901–1930* |  |  |  |  |  |
| Full study area | 0.01 (-0.02, 0.05)<sup>†</sup> | -0.10 | -0.04 | – | -0.20 |
| Core range | 0.01 (-0.01, 0.06)<sup>†</sup> | -0.26 | n.s. | – | -0.18 |
| Outside core range | 0.00 (-0.02, 0.04)<sup>†</sup> | 0.09 | -0.04 | – | -0.26 |

\*Spearman's $\rho$ reported if $p < 10^{-3}$ or better, n.s. otherwise
<sup>†</sup> greater than zero, Wilcoxon sign-rank test $p < 10^{-6}$

**Fig. 4.**
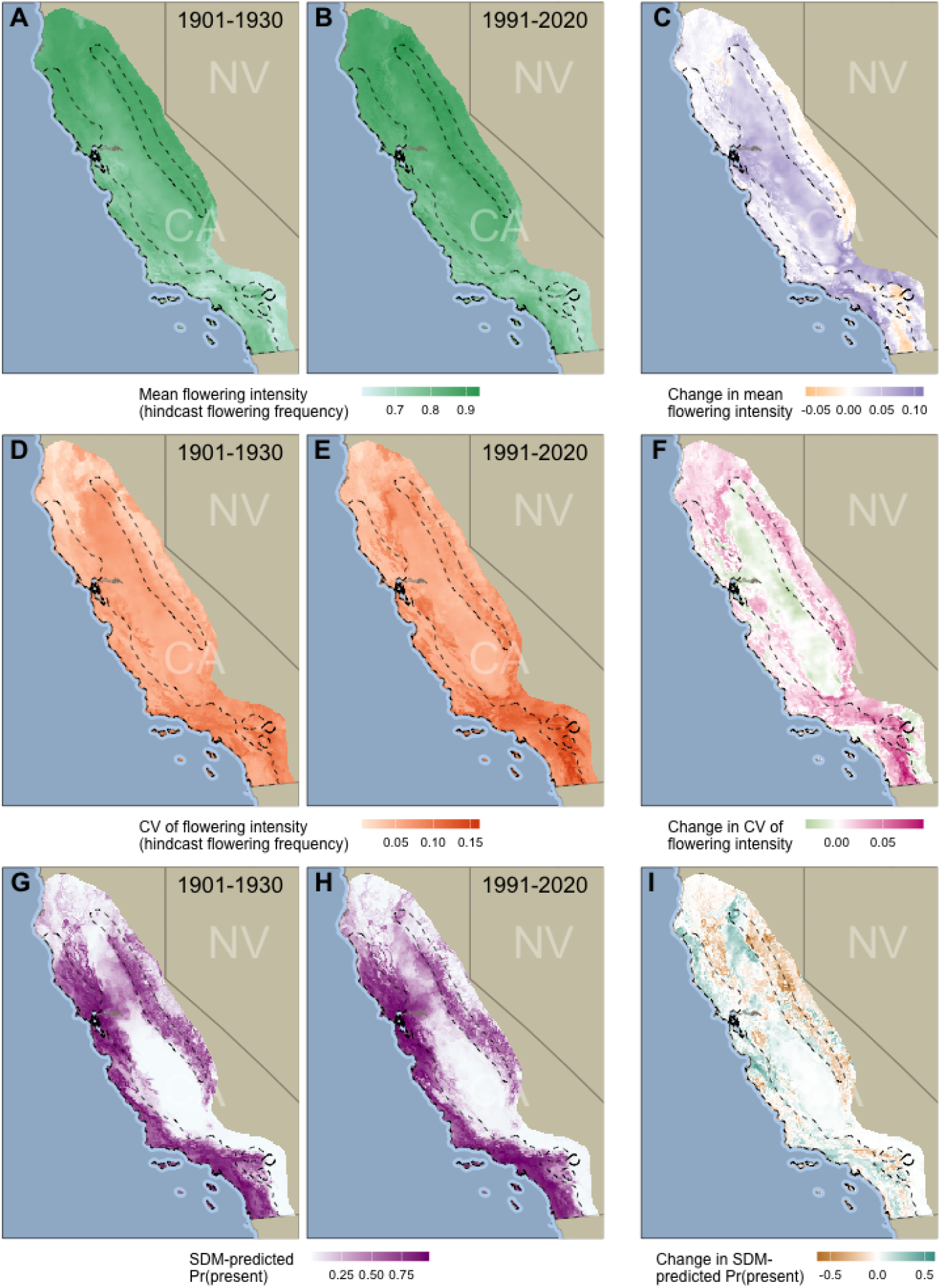
Hindcast changes in toyon flowering from the early 20th century to recent decades, compared to predicted probability of occurrence from a static SDM. (A) Mean hindcast flowering intensity over the years 1901–1930; (B) Mean flowering intensity over 1991–2020; (C) Change in mean flowering intensity from the early period to the recent period; (D) CV of flowering intensity over 1901–1930; (E) CV of flowering intensity over 1991–2020; (F) Change in CV of flowering intensity from the early period to the recent period. (G) SDM-predicted probability of toyon presence based on climate averages for 1901–1930; (H) SDM-predicted probability of occurrence based on climate averages for 1991–2020; and (I) change in SDM-predicted probability of occurrence from the early period to the recent period. In all panels, dashed outlines indicate the toyon core range.

To validate the hindcast predictions, we compared hindcast flowering intensity in given years and locations to flowering intensity estimated from surveys compiled by USA-NPN, representing flowering years 2010–2024 at 197 unique sites (USA National Phenology Network, 2025). The hind-cast prediction of flowering intensity was significantly and positively correlated with the flowering intensity derived from USA-NPN survey data (Spearman’s *ρ* = 0.23, *p* < 10^−5^), consistent with the expectation that the hindcast accurately reflected flowering intensity in the years and locations represented in that independent data set.

### Hindcast changes in flowering intensity

The hindcast projects that flowering intensity increased slightly in 1991–2020 compared to 1901–1930 (Fig. 4A–C; Table 1): the median change in mean flowering intensity across the whole study area was 0.02 (95% density from −0.02 to 0.08), with similar changes inside and outside of the core range (all significantly greater than 0; Wilcoxon sign-rank test *p* < 10^−6^). Increases in flowering intensity were stronger in the Los Angeles Basin and the Central Valley, and the most striking decreases were in inland Southern California (Fig. 4C). Variation in flowering intensity also increased slightly (Fig. 4D–F; Table 1): median change in the CV of flowering intensity was 0.01 (−0.02 to 0.05), with similar changes inside and outside of the core range (all significantly greater than 0; Wilcoxon sign-rank test *p* < 10^−6^). The biggest increases in CV of flowering intensity were in the southern part of the range, but variation increased in higher elevation regions from the north to the south (Fig. 4F).

These changes in hindcast flowering intensity follow from changes in the weather predictors of flowering used by our model. Comparing changes in the predictors to changes in hindcast flowering activity, we found that increases in mean flowering intensity are most strongly associated with changes in precipitation (Fig. 5A): regions with larger increases in mean flowering intensity have seen greater increases in precipitation in the winter of the flowering year (PPT Y0Q1) and in the summer before flowering (PPT Y1Q3), and associated declines in VPD in the winter of the flowering year (VPDmin Y0Q1); relationships with changes in the three other predictors were shallower. In contrast, greater increases in the CV of flowering intensity were in regions with declining precipitation, rising minimum temperatures, and associated increases in VPD (Fig. 5). These patterns are consistent with increasing mean flowering intensity in regions that have become somewhat more hospitable for toyon, and increasing inter-annual variation in flowering intensity in regions that have become more stressful since the early 20th century.

**Fig. 5.**
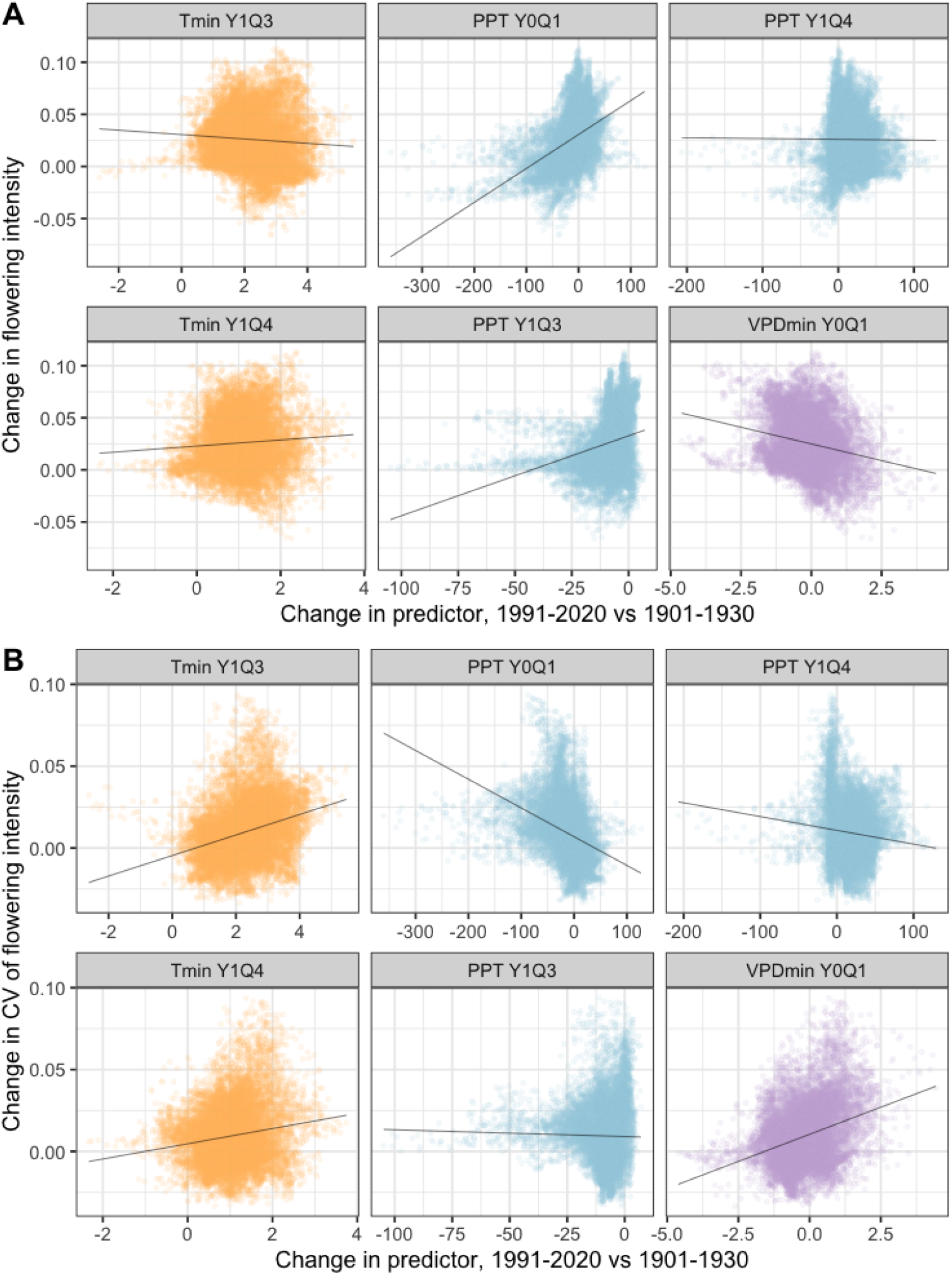
Change hindcast flowering compared to change in mean predictor values, 1901–1930 versus 1991–2020. For each panel, the y-axis value gives the change in (A) mean or (B) CV of hindcast flowering intensity for each raster cell, and the x-axis gives the change in the mean value of the corresponding predictor. Lines are linear regressions. Predictor names and coloring follow Fig. 3B.

### Hindcast flowering in *a priori* and projected climate refugia

If flowering intensity is a component of toyon population health, we would expect that changing global climates should result in stable to increasing flowering intensity at higher latitudes and higher elevations, and in regions identified by a presence-based SDM as gaining suitability for toyon since the early 20th century. Flowering intensity was strongly and positively correlated with latitude in both the early (1901–1930) and recent (1991–2020) periods, whether considered across the full study area, within the core range, or outside the core range (Table 1; Spearman’s *ρ* ≥ 0.81, and *p* < 10^−3^ in all cases). However, across the full study area, change in mean flowering intensity was significantly negatively correlated with latitude (*ρ* = − 0.22, *p* < 10^−3^). This effect was mostly driven by regions outside the core range of toyon, as within the core range the correlation was much smaller, though still significantly different from zero (*ρ* = − 0.34 outside the core range, *ρ* = − 0.08 inside, *p* < 10^−3^ in both cases).

Variation in flowering was strongly negatively correlated with latitude (i.e., CV of flowering intensity was lower farther north) in both time periods, and both inside and outside the core range (Table 1; Spearman’s *ρ* ≤ 0.52, and *p* < 10^−3^ in all cases). In parallel with change in mean flowering intensity, the correlation between latitude and change in the CV of flowering intensity differed inside and outside of toyon’s core range: inside the core range, the change in CV of flowering intensity was strongly negative (*ρ* = − 0.26, *p* < 10^−3^); outside, this correlation was weakly but significantly positive (*ρ* = 0.09, *p* < 10^−3^). These results — comparatively little change in the relationship between flowering and latitude, and a significant reduction in the relationship between variation and latitude — are consistent with conditions favoring flowering remaining more stable in the core range.

Hindcast mean and variation of flowering intensity were more weakly, in many cases non-significantly, correlated with elevation (Table 1). Mean hindcast flowering intensity for both the early and recent periods were weakly positively correlated with elevation within the core range (*ρ* = 0.07 in the early period, *ρ* = 0.05 in the recent, *p* < 10^−3^ in both cases), but had no significant correlation outside the core range; change in mean flowering intensity was not significantly correlated with elevation inside the core range, but it was significantly greater at higher elevations outside the core range (*ρ* = 0.07, *p* < 10^−3^). Year-to-year variation in flowering intensity was negatively correlated with elevation, both across time periods and regions (*ρ* ≤ − 0.08 and *p* < 10^−3^ in all cases), while change in the CV of flowering intensity was not significantly correlated with elevation in the core range, and weakly negatively correlated with it outside the core range (*ρ* = −0.04, *p* < 10^−3^).

Finally, hindcast flowering intensity did not show a consistent relationship to probabilities of toyon occurrence (*Pr*(*present*)) predicted by the static SDM, but did align with change in the predicted probability of occurrence (Table 1). Within the core range, hindcast flowering intensity was significantly negatively correlated with SDM-predicted probability of toyon presence, in both the early and recent periods (*ρ* = − 0.19 for the early period, *ρ* = − 0.20 for the recent period, *p* < 10^−3^ for both). Outside the core range, this correlation was reversed (*ρ* = 0.36 for the early period, *ρ* = 0.19 for the recent period, *p* < 10^−3^ for both). Change in mean hindcast flowering intensity was positively correlated with change in SDM-predicted probability of occurrence both inside and outside the core range, however (*ρ* = 0.09 inside the core range, *ρ* = 0.22 outside, *p* < 10^−3^ for both).

Similarly, the correlation between variation in flowering intensity and SDM-predicted probability of occurrence varied across regions or time periods, but there was a more consistent relationship between change in the CV of flowering intensity and change in SDM-predicted probability of occurrence. In the core range, CV of flowering intensity was positively correlated with probability of occurrence in the early period (i.e., greater variation in regions with higher probability of occurrence; *ρ* = 0.23, *p* < 10^−3^), but had no significant correlation in the recent period. Outside the core range, there was a negative correlation between CV of flowering intensity and probability of occurrence in the early period (*ρ* = − 0.07, *p* < 10^−3^), which became a positive correlation in the recent period (*ρ* = 0.12, *p* < 10^−3^). However, change in the CV of flowering intensity was negatively correlated with change in the probability of occurrence across the study area (*ρ* = − 0.18 inside the core range, *ρ* = − 0.26 outside, *p* < 10^−3^ for both).

Overall, SDM-predicted probability of occurrence was not clearly aligned with predicted flowering activity at the smaller spatial and temporal scales. Nevertheless, SDM-predicted increases in the probability of toyon occurrence over a 120-year period did align with TARDIS predictions of increasing flowering intensity, and decreasing inter-annual variation in flowering.

## Discussion

Species’ geographic distributions arise from the interaction of their environmental tolerances with their dispersal capacity and the range of habitat conditions they encounter. The TARDIS method, modeling reproductive activity over time in relation to annual weather variation, may help to parse the relationship between where a species can maintain a self-sustaining population (i.e., its fundamental niche) and its realized geographic distribution (Holt, 2009). Our analysis of toyon flowering activity reveals how this ecologically and culturally important species has experienced variation in habitat quality across space and over 125 years of global change, and suggests that most toyon populations remain demographically healthy, consistent with a stable geographic distribution as climate changes.

Our continuous-response BART model recovers biologically realistic weather predictors of flowering intensity (Fig. 3), and predicts flowering activity in alignment with independent observations of toyon flowering recorded by USA-NPN (USA National Phenology Network, 2025). Using the continuous-response model to hindcast toyon flowering activity for each year from 1900 to 2024, we find that conditions favoring flowering have been stable to slightly increasing across much of toyon’s range (Fig. 4C). This trend is somewhat more positive at higher elevations, but not at higher latitudes, and it aligns with changes in the probability of toyon occurrence predicted by a conventional SDM (Fig. 4C,I; Table 1). We also find that inter-annual variation in predicted flowering intensity declines at higher latitudes and higher elevations, and that regions with increasing SDM-predicted probability of occurrence also have decreasing inter-annual variation in conditions favoring flowering (Fig. 4F,I; Table 1). Changes in the mean and CV of flowering intensity predicted in our analysis reflect changes in temperature and precipitation from the early 20th century to recent decades (Fig. 5). These results reinforce prior work indicating that toyon’s distribution is not likely to shift substantially with ongoing climate change (Riordan et al., 2018), and suggest how key population processes (i.e., flowering and fruiting) contribute to the presence of this foundational species on the landscape.

### Matching model form to life history

TARDIS was first developed to study a masting species, which invests in mass-flowering events separated by one or more years without substantial flowering (Yoder et al., 2024). Because toyon varies in annual flowering intensity but generally has at least some investment in flowering and fruit production every year (Fig. 1D), we considered whether a different model structure was appropriate: treating flowering activity as a continuous response — frequency of records indicating flowering — rather than the binary presence/absence response that Yoder *et al* (2024) used to model masting. We trained and compared binary- and continuous-response BART models of toyon flowering, and found that although both model forms identified overlapping sets of top predictors with qualitatively similar partial effects (Figs 2, 3), the continuous-response model explained notably more variation in flowering activity than the binary-response model (RMSE 0.35 for continuous-response vs 0.59 for binary-response). This comparison thus highlights the importance of considering biological realism in model development — both *a priori*, noting that toyon flowering better conforms to a continuous than a binary response, and in consideration of accuracy seen for the trained model. Overall, this study provides a proof of concept for TARDIS with a model structure that will likely apply to a broader range of perennial plant species than the binary-response model demonstrated by Yoder *et al* (2024).

### Modeling species distributions as more than presence

A challenge for inferring species’ habitat needs from the range of environments in which they occur is that species may be absent from places where they could maintain healthy populations (because of dispersal limitation), and present in places that do not support net population growth (because conditions have changed since long-lived individuals established, or because of immigration from more suitable locations; Anderson 2013; Holt 2009; “habitat debt” per Radomski 2025). TARDIS aims to improve over presence-based SDMs by modeling annualized reproductive activity, rather than a species’ presence on the landscape. To see whether TARDIS accomplishes this aim, in this study we make an explicit comparison between TARDIS-predicted flowering activity, and the probability of toyon presence (habitat suitability) predicted by a presence-based SDM.

We find a complex relationship between flowering intensity and presence, varying with geographic and temporal scope (Table 1). Across the full study area, predicted flowering intensity for 1991–2020 is weakly negatively correlated with predicted probability of toyon occurrence (*ρ* = − 0.05, *p* < 10^−3^), which masks strongly contrasting correlations inside the core range (*ρ* = − 0.20, *p* < 10^−3^) and outside it (*ρ* = 0.19, *p* < 10^−3^). This is potentially consistent with toyon populations in the core range experiencing conditions that have become less suitable since the long-lived plants established, as “trailing edge” populations, while regions beyond the core range have become better suited for toyon recruitment (Kroiss and Hille Ris Lambers, 2015; Walck et al., 2011). In 1901–1930, prior to the warming trends of the 20th century, we do see a weak positive correlation between predicted flowering intensity and probability of occurrence across the full study region (*ρ* = 0.04, *p* < 10^−3^), but there is still a negative correlation within the core range *ρ* = − 0.19, *p* < 10^−3^), almost identical to that seen for the core range in the recent period (*ρ* = − 0.20, *p* < 10^−3^). Moreover, there is a consistent positive correlation between *change* in flowering intensity and probability of presence from the early period to the recent one — regions that have seen greater increases in probability of toyon occurrence in 1991–2020 compared to 1901– 1930 have also seen greater increases in predicted flowering intensity (Table 1; *ρ* = 0.16, across the full study region, *ρ* = 0.09 in the core range, *ρ* = 0.22 outside the core range, *p* < 10−3 in all cases).

This complexity may be explained by the fact that these correlations capture two different sources of variation: geographic variation in flowering intensity within the early or recent time periods is not necessarily aligned with SDM-predicted probability of presence — but changes in flowering intensity and probability of presence occur in the same direction across our study area. This suggests that, for the 125-year time frame of climate change captured by our study, the presence-based SDM provides a good proxy for population processes (flowering) that contribute to presence, even though presence and demography are not well aligned over shorter time frames.

It is also possible our results reflect range limits defined not by average environmental conditions but by inter-annual variation in conditions. Theory of range limits has long recognized that increasing variation in habitat suitability, and thereby demographic status, may define range edges (Birch 1957; Curnutt et al. 1996; Richards 1961; reviewed by Pironon et al. 2017). Classic metapopulation models of occupancy across an environmental gradient show increasing variation in population status towards the edge of the realized distribution, created by higher population turnover in less-suitable conditions (Holt and Keitt, 2000; Lennon et al., 1997), and evolutionary modeling shows stable range limits can be created by increasing genetic load specifically as a result of greater environmental variation at the range edge (Benning et al., 2022). We find a higher CV of modeled annual flowering intensity at lower latitudes and lower elevations, consistent with more marginal conditions for toyon, in both the early and recent periods (Table 1). We also find a negative correlation between change in the CV of modeled flowering intensity and change in suitability predicted by the presence-based SDM, consistent with the hypothesis that variation in population status defines the edge of habitat suitability for toyon. If that is the case, our finding that the CV of flowering intensity has slightly increased from the early period to the recent period would suggest slightly declining habitat suitability for toyon, even as mean flowering intensity remained stable to increasing (Table 1).

### Toyon’s fundamental niche and geographic distribution in a changing world

The California Floristic Province (CFP) is a global hotspot for plant biodiversity, home to unique communities assembled through the interactions of complex geological substrates and a Mediterranean climate (Millar, 2012) — and global change is already reshaping this diversity (Crimmins et al., 2011; Harrison et al., 2024). Predicting these changes and protecting the region’s endemic diversity is an on-going challenge, and our analysis here demonstrates a general approach for improved predictions of plants’ responses to changing temperature and precipitation through the specific case of a foundational species in CFP chaparral and woodland communities.

Our results are consistent with prior work finding the geographic distribution of optimal conditions for toyon should be largely stable under recent and future climate change, and that shifting precipitation regimes may have the biggest impact on CFP biodiversity under global change. A prominent study by Crimmins *et al*. (2011) drew on historic surveys of 46 California plant species to find that most had optimal climate conditions shifting not uphill but downhill, following changing water balance rather than temperature alone — and specifically that toyon’s optimal range moved 53 meters downhill. Our analysis of change in conditions favoring toyon flowering largely aligns with these results: we find that change in flowering intensity is substantially driven by change in precipitation regimes (Fig. 5), and that it is not significantly correlated with elevation in toyon’s core range (Table 1). Toyon is also among the species considered in a SDM study by Riordan *et al*. (2018) projecting changes to southern California shrubland communities under future climate scenarios for 2040–2069, which found that, averaged across scenarios, 99% of toyon’s current habitat should remain suitable into the mid-21st century. Comparing our results to these prior studies of toyon’s realized distribution supports the idea that toyon is likely to persist in its current range is supported by conditions that maintain a key component of population health, annual reproductive activity.

These studies and ours further suggest that future risks to toyon and many (though by no means all; Riordan et al. 2018) CFP endemics arise not so much from direct effects of global climate change, but from secondary effects such as changing fire regimes and species invasions, and from concurrent habitat loss to development (Goss et al., 2020; Lawson et al., 2010; Lenihan et al., 2003; Riordan and Rundel, 2014; Rose et al., 2023; Williams et al., 2019). Our hindcast finds increasing year-to-year variation in flowering intensity (Fig. 4F) driven by changing precipitation patterns (Fig. 5), and previous studies have also found that increasing annual variation in precipitation — swings from drought to extreme rainfall years — is a substantial source of climate change risk to CFP communities (Berg and Hall, 2015; Harrison et al., 2024). Thus, although TARDIS finds conditions supporting toyon populations’ reproductive activity have been stable to increasing over our study period, our results are consistent with complex, interacting risks for the species as global change continues.

### Understanding population status via crowd-sourced natural history observations

This project joins a growing body of research making use of casually collected natural history observations to address questions in ecology. Volunteer contributors to “citizen science” or “participatory science” projects (Cooper et al., 2021; Ellwood et al., 2023) can often provide broader geographic, temporal, and taxonomic coverage than directed efforts by working scientists (Amano et al., 2016; Dickinson et al., 2010; Panter et al., 2020). Crowdsourcing can produce data in alignment with expert-collected observations (Aceves-Bueno et al. 2017; Callaghan et al. 2020, but see Tiago et al. 2017), especially with appropriate quality controls (Kosmala et al., 2016; Panter et al., 2020). This kind of approach has already provided valuable data for species distribution modeling (Gaier and Resasco, 2023; Tiago et al., 2017); quantifying and monitoring biodiversity (Fink et al., 2023; Pernat et al., 2021; Wilson et al., 2020) — and for studying the timing and intensity of plants’ reproductive activity under changing climates (Brenskelle et al., 2021; Chung et al., 2011; Yoder et al., 2024).

For toyon, iNaturalist contributors provided thousands of records of flowering and fruiting status that were usable for model training (Fig. 1). Species identity for these records could be readily validated from the attached images — *Heteromeles* is a monospecific genus, and toyon’s leaf morphology is distinguishable from that of a few co-occurring species with superficially similar fruit or flowers. Some potential sources of bias are obvious in the data all the same: the growth of the iNaturalist contributor community means that records are heavily skewed towards recent years (Fig. 1D), and records are spatially clustered around human population centers in the San Francisco Bay area, the Los Angeles basin, and around San Diego (Fig. 1D). Nevertheless we do not see a temporal trend in the frequency of records indicating flowering, the key response variable in our models, and despite spatial clustering our training data samples the geographic breadth of the species’ range. These criteria give us some confidence that temporal and spatial heterogeneity in our crowdsourced sample are not a source of systematic error (Yoder et al., 2024).

### Future directions for TARDIS

Our modifications to the analysis pipeline of Yoder *et al*. (2024) let us study species whose life histories are not defined by periodic masting events. This substantially broadens the possible applications of TARDIS, but it does not accommodate the full diversity of plant life histories. Annual species, which are only apparent on the landscape when reproductive, do not offer the same opportunity to train models on “true absence” observations that we can obtain for long-lived perennials like toyon. Then, too, the reproductive activity modeled by TARDIS is only one component of population demographic health — population growth or decline is also determined by establishment and survival of seedlings, and mortality of reproductive individuals. Although focusing on the life-history stage that is most accessible via crowdsourced data is a key advantage of our approach, it would be interesting to adapt TARDIS to model more elements of the life cycle. This might amount to recreating more complex mechanistic SDMs (Merow et al., 2014a,*b*), but a more comprehensive TARDIS could identify how specific life history stages contribute to population health and persistence.

## Conclusions

Understanding how species’ geographic distributions are shaped by environmental variation over time and space is both a fundamental task of ecology, and an important part of managing natural communities’ response to on-going climate change (Radomski, 2025; Thomas, 2010). Adapting the TARDIS pipeline of Yoder *et al*. (2024) to study flowering in toyon populations reveals how changing winter precipitation and temperatures have likely shifted the intensity of toyon flowering in many parts of its range (Fig. 5). Trends in our hindcast of toyon flowering from 1900 to 2024 suggest populations of this ecologically important species remain healthy, consistent with predictions from a presence-based distribution model that finds relatively little change in the geographic distribution of suitable habitat for toyon over the same period. At the same time, our analyses indicate toyon may have little opportunity to migrate north under continuing climate change, even as it faces threats beyond changing rainfall and temperature regimes. The endemic diversity of the California Floristic Province faces multiple overlapping threats in a warming world, and while we find reason to believe in the resiliency of toyon populations, it would be better for this iconic species, and the unique communities in which it grows, if that resiliency were not tested further.

## Acknowledgments

We wish to thank editors Jeremy Rentsch, Elizabeth Stacy, Caitlin Mackenzie, Jennifer Boyd, and Vivian Negron-Ortiz for the invitation to contribute to this Special Issue on Plant Resilience and Conservation for a Changing Climate. Support was provided by the U.S. National Science Foundation (DEB 2001180). Data were provided by the USA National Phenology Network and the many participants who contribute to its *Nature’s Notebook* program, and by the iNaturalist contributor community, without whom this project would not be possible. Thanks are also due to two anonymous reviewers for their constructive comments, to Colin J. Carlson for critical feedback on model development, and to Richard Rogers and Oscar Hammerstein II for making raindrops on the flowers of a rosaceous shrub one of our favorite things.

## Author contributions

DD and JBY conceived and planned the project; DD annotated crowdsourced records and managed data compilation; JBY developed the analysis; DD and JBY conducted analysis and drafted the manuscript.

## Data availability

Code for acquisition and management of training data, and for running all analyses, is available on GitHub: github.com/JBYoderLab/flower_prediction_toyon

